# Conditioned Protein Structure Prediction

**DOI:** 10.1101/2023.09.24.559171

**Authors:** Tengyu Xie, Zilin Song, Jing Huang

**Affiliations:** Westlake University

## Abstract

Deep learning based protein structure prediction has facilitated major breakthroughs in biological sciences. However, current methods struggle with alternative conformation prediction and offer limited integration of expert knowledge on protein dynamics. We introduce AFEXplorer, a generic approach that tailors AlphaFold predictions to user-defined constraints in coarse coordinate spaces by optimizing embedding features. Its effectiveness in generating functional protein conformations in accordance with predefined conditions were demonstrated through comprehensive examples. AFEXplorer serves as a versatile platform for conditioned protein structure prediction, bridging the gap between automated models and domain-specific insights.

## 1 Introduction

Contemporary structural biology examines not only the architecture of biomacromolecules but also discerns their functional implications, which are of great scientific and therapeutic importance. In recent years, the field has witnessed transformative advancements in computational structural biology empowered by deep learning (DL) approaches such as `AlphaFold2` (AF) and comparable methods [1, 2, 3, 4, 5, 6, 7]. These tools have leveraged protein structures determined by laborious experiments such as X-ray crystallography, NMR spectroscopy, or cryogenic electron microscopy, facilitating the widely accepted interim breakthroughs in the highly accurate prediction of *static* protein structures [8]. However, physiologically relevant protein motions, especially in functional response to external perturbations or interactions, are highly dynamic.

Despite successful attempts have been made to capture the ensemble and dynamic nature of protein conformations, effective applications of AF on modern protein science research have yet to be fully realized due to the lack of interoperability between these data-driven models and human intelligence [9]. Current DL models could not generate protein structures with specific local or global structural features, as the domain knowledge from human experts on specific conformational descriptors that distinguish different protein functional states is not embedded [10]. For example, AF lacks the capability to specifically generate the conformational structure of a membrane transporter in its outward-facing state. Such discrepancy impedes the robust and productive applications of AF and related methods in crucial research practices such as the developments of novel therapeutic agents or biosynthetic pathways.

Here, we present `AFEXplorer` (AFEX), a versatile approach that offers highly accessible user-conditioned protein structure prediction with AF. In brief, AFEX serves as a surrogate protein structure synthesizer that learns the optimal feature representation for AF to generate protein conformations subjected to spatial constraints in a coarse coordinate space.

## 2 Related Works

### 2.1 Physics-based modeling

The conformational change of proteins associated with the transition between functional states is often a rare event in protein dynamics. Theoretical understanding of rare event sampling predominantly comes from physics-based modeling. Among various rare event enhanced sampling methods, Targeted Molecular Dynamics (TMD) [11] is especially useful for directing the time evolution of protein conformations towards specific configurations. Basically, TMD introduces to the *N* -body subsystem an extra penalty potential, U_bias_, on the instantaneous coordinates **x**(*t*):

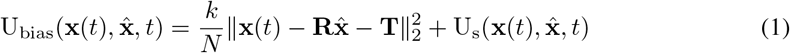

where *k* is the force constant, 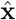 represents the target subsystem configuration, U_s_ is a time-scheduled smoothing potential, and the alignment operators 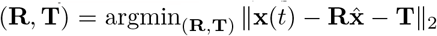. As long as the Boltzmann statistics apply, time evolution of the system would approach the targeted configuration under this bias potential. A closely related method for studying protein unfolding dynamics is Steered Molecular Dynamics (SMD), which exerts a pulling force of fictitious origin on the terminal region to distort the folded structure [12]. In this approach, equilibrium thermodynamic observables can be retrieved via the Jarzynski equation in the case of irreversible pulling [13].

### 2.2 Generating alternative structures with AF

Interest in exploring the conformational ensemble learned by AF has catalyzed an evolving area of research [14, 15]. Indeed, AF or equivalent models can be viewed as a black-box function that maps the multiple sequence alignment (MSA) featurization of amino acid sequences to the corresponding protein spatial coordinates. One could thus characterize multiple protein conformations by feeding AF with randomly perturbed or specifically curated feature representations. For example, Heo and Feig proposed to substitute the default template database with a curated set of GPCR homologues in the active state, aiming to mitigate AF’s bias toward predicting inactive GPCR conformations [16]. Similarly, a strategy that combines MSA subsampling [17] and template manipulation has been exploited to model the functional conformations of kinases and GPCRs [18].

Different ways to subsample the raw MSA inputs have been proposed [17, 19, 20]. Del Alamo et al. adopted stochastic sampling to reduce the depths of MSAs, aiming to drive AF to produce alternative structures with some likelihood of differing from the default AF predictions [17]. Wayment-Steele et al. developed the `AF-Cluster` method, which clusters MSAs to maximize the coevolutionary decoupling between distinct folds and use these fold-aware MSA clusters to generate alternative structures via AF [19]. A recent benchmark on 16 membrane proteins indicates limited success of these two methods in alternative conformation predictions [21]. Zhang et al. developed an MSA generative model that, by feeding random noises, can generate various MSA sets with the potential to drive AF to produce alternative conformations [22]. Moreover, the `AFsample` approach proposed by Wallner uniquely sets the dropout layers in AF-Multimer to training mode during inference; models stochasticity induced by random dropouts in the internal AF latent representation naturally lead to diverse protein complex conformation predictions [23]. Recently, Bryant and Noe introduced `AFProfile`, which uses AF-Multimer confidence scores as the objective to learn an offset bias that denoises the input MSA for improving AF-Multimer predictions [24].

## 3 Method

We redefine the conditioned structure prediction as an optimization problem, where the objective is to identify the optimal MSA feature that forces AF to generate the structures that align with our *apriori* knowledge on specific states (Figure 1).

**Figure 1.**
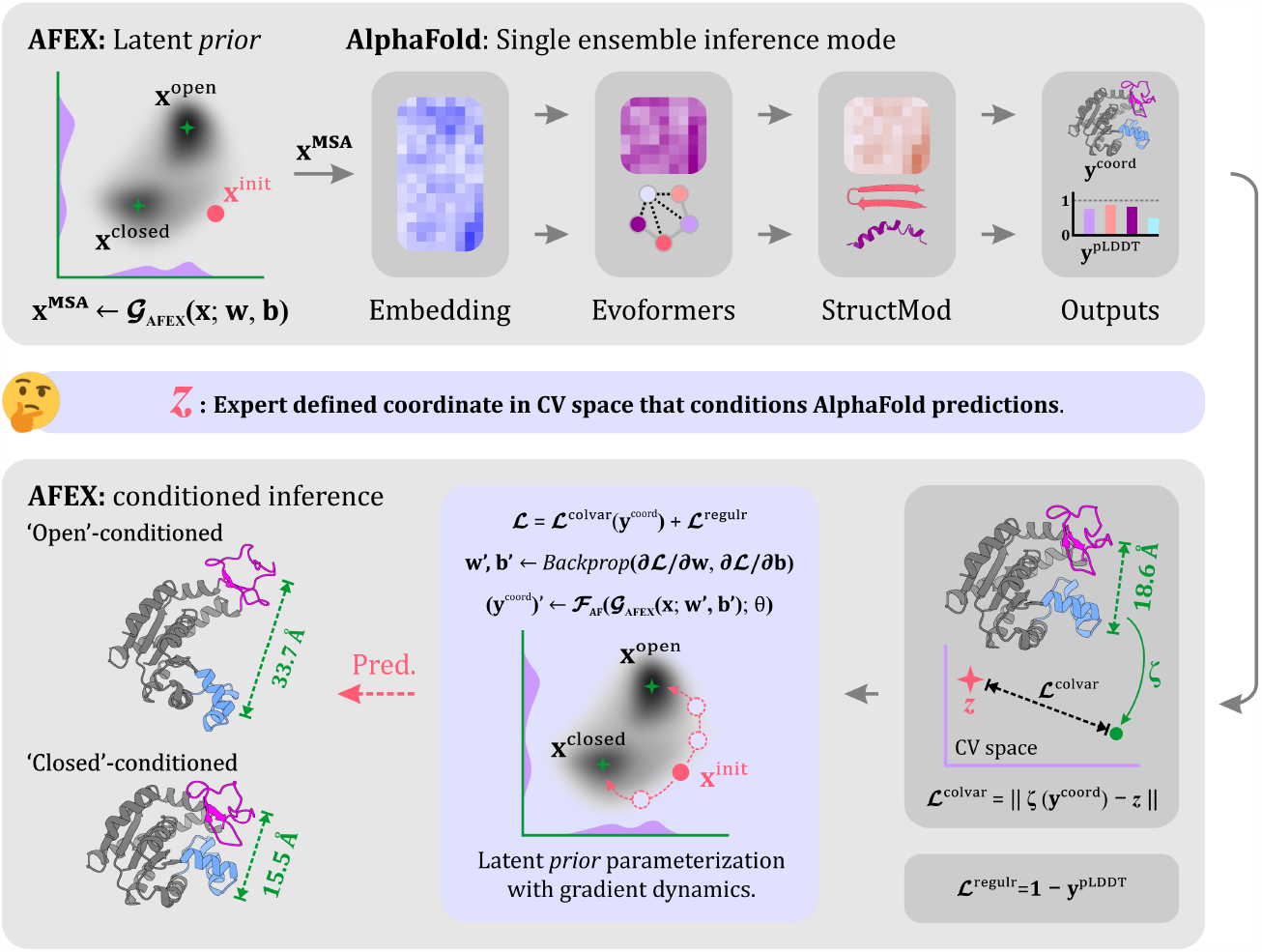
The AFEX protocol is designed to identify MSA features tailored to specific conformational states. It leverages domain knowledge, encoded as a loss function, to optimize features that guide AF in generating the targeted state. The AdK protein serves as an example to demonstrate AFEX’s capability in generating both its open and closed states.

### 3.1 Algorithm

We begin with the AF inference function, ℱ_**AF**_, whose input could be generally categorized into features of the predicting target (**x**^targ^), the MSA profiles (**x**^MSA^), and the template profiles (**x**^temp^):

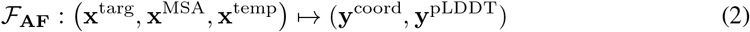

where **y**^coord^ and **y**^pLDDT^ denotes the predicted Cartesian coordinates and the per-residue confidence of the prediction. Let us introduce the *a prior* knowledge from human experts that a reduced representation *ξ* : **y**^coord^ ↦*z*^colvar^ could *reasonably* capture the functional evolution of the target protein conformations and we are interested in a specific state, 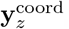, characterized by the Dirac’s *δ* function 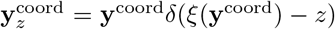. To find 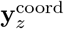through ℱ_**AF**_, we look for 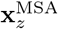as we keep the target features **x**^targ^ fixed and drop the template inputs to *ℱ*_**AF**_:

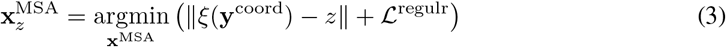

where *II & II* denotes a generic distance measure that defines the collective variable (CV) loss *ℒ*colvar, and *ℒ*^regulr^ represents the regularization loss. We find that 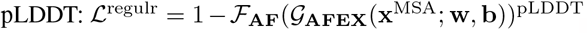 can be parametrized via a generator, *𝒢*_**AFEX**_:

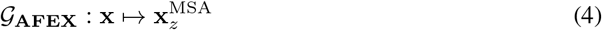

where **x** denotes *some* latent prior. This concludes the AFEX framework with Eqs. (2), (3), and (4). While extensions for learning optimal feature representations other than MSAs (e.g., templates) are feasible, we note that many previous methods for diversifying AF predictions based on feature perturbations can be viewed as special cases of AFEX with specific implementations of *𝒢*_**AFEX**_.

Although more complex differentiable architectures can be parametrized as the latent feature generator, in the present work, we simply implement 𝒢_**AFEX**_ as linear activation applied selectively to the `cluster_profiles` of the MSA features:

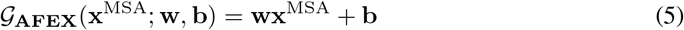

Based on such a linear mapping, the inference graph

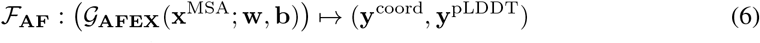

naturally casts to the optimization problem:

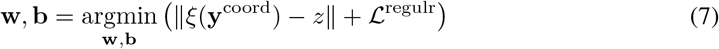

with the regularization loss defined using pLDDT: *ℒ*^regulr^ = 1 *−*ℱ_**AF**_(𝒢_**AFEX**_(**x**^MSA^; **w, b**))^pLDDT^.

### 3.2 Implementation details

The implementation of AFEX requires no modifications to the AF source codes (as per Eq. (6)). The AF runtime is kept in inference mode, with a single MSA feature ensemble. The CV losses used for each tested system are detailed in their respective Results sections. We parametrized each linear AFEX learner with the Adam optimizer [25]. The implementation leverages the `Optax` library within the `JAX` ecosystem [26]. AFEX is publicly available at: https://github.com/JingHuangLab/AFEXplorer.

## 4 Results

### 4.1 AFEX generates AdK open and closed states

Adenylate kinase (AdK) is a classic system for studying large-scale conformational changes [27]. We define the CV as the distance (*D*) between the C*α* atoms of A37 in the AMP lid and R124 in the ATP lid (E. *coli* AdK sequence). The corresponding loss function is defined using the sigmoid function *σ*(*x*) = 1*/*(1 + exp(*−x*)) as 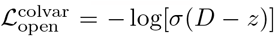 and 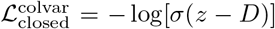 to 25 Å and ℒ^regulr^ is added to 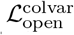 or 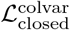 to formulate the final loss function of AFEX. A 500-step optimization of **w** and **b** drives the AF prediction towards the open conformation (*D* = 33.7 Å) and the closed conformation (*D* = 15.5 Å), compared to the initial *D* value of 18.6 Å (Figure 1). The overlap with experimental AdK structures 4AKE (*D* = 33.6 Å) and 1AKE (*D* = 16.1 Å) confirms that highly accurate structure predictions for these two functional states were obtained. The C*α* RMSDs are 1.60 Å for the open and 1.83 Å for the closed conformation, respectively.

### 4.2 AFEX generates DFG-in/out conformations for all human kinases

Kinases are important drug targets due to their central role in regulating cellular signaling pathways. They are highly dynamic, capable of adopting various conformations such as the DFG-in and DFG-out states [28]. The DFG-out conformation is important because it creates a binding pocket that facilitates the design of type-II inhibitors, with imatinib (Gleevec) serving as a prominent example. Capturing the DFG-out conformation for a kinase remains a challenge even with the advent of AF, as AF exhibits a strong bias towards the DFG-in state [29, 30]. Modi and Dunbrack identified two CVs, *D*_1_ and *D*_2_, for distinguishing DFG-in and DFG-out states. *D*_1_ is the distance between the C*ζ* atom of the Phe residue in the DFG motif and the C*α* atom of the fourth residue following the conserved Glu in the C-helix, while *D*_2_ is the distance between the Phe C*ζ* atom and the C*α* atom of the conserved Lys in the *β*3 strand [28]. DFG-in is indicated by *D*_1_ *≤* 11 Å and *D*_2_ *≥* 11 Å, while *D*_1_ *>* 11 Å and *D*_2_ *≤* 14 correspond to DFG-out.

Accordingly, we perform AFEX calculations using the CV loss defined as 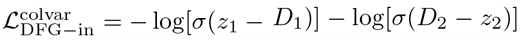 for DFG-in and 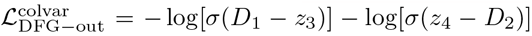 for DFG-out, with *z*_1_ = *z*_2_ = *z*_3_ = 11 Å and *z*_4_ = 14 Å. Combined with ℒ^regulr^, we generated specific DFG-in and DFG-out conformations for 453 human kinases. As shown in Fig. 2, 99.3% of DFG-in and 99.1% of DFG-out conditioned predictions were successful. In contrast, AF generates predominantly DFG-in states with very narrow distributions (Fig. 2d). For a specific kinase, the two conformations were generally similar, differing mainly in the local structural arrangement we conditioned on. Associated variations in adjacent structural elements, such as the activation loop, were also observed (Fig. 2e). While AFEX expands the accessible conformational space of human kinases, the distributions of *D*_1_ and *D*_2_ differ from those in the PDB as plotted in Fig. 2c using 7,978 experimental kinase structures. Further refinement may be achievable through more sophisticated definition of CV loss.

**Figure 2.**
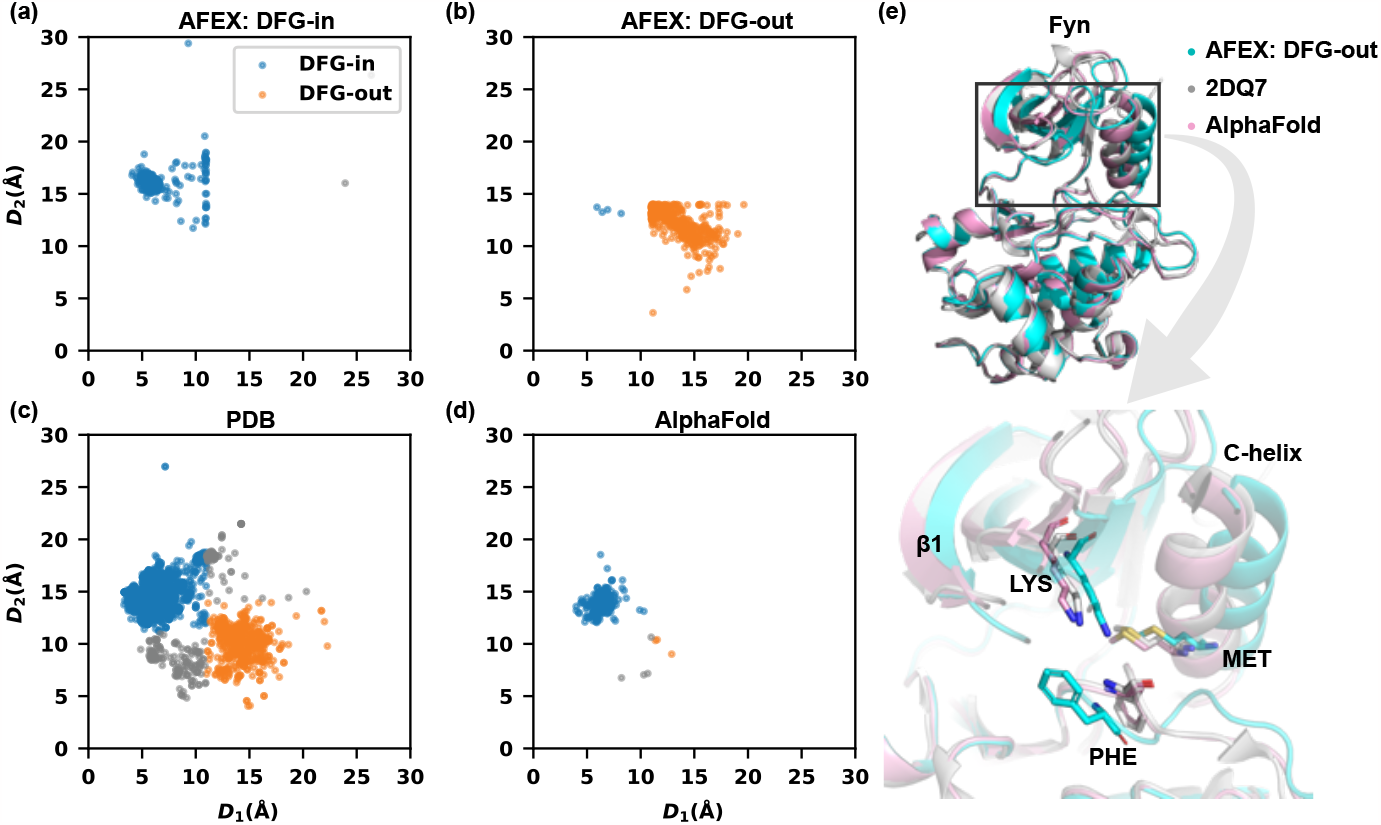
AFEX successfully generated alternative states (DFG-in and DFG-out) for human kinases. The Fyn kinase is shown as an example.

**Figure 3.**
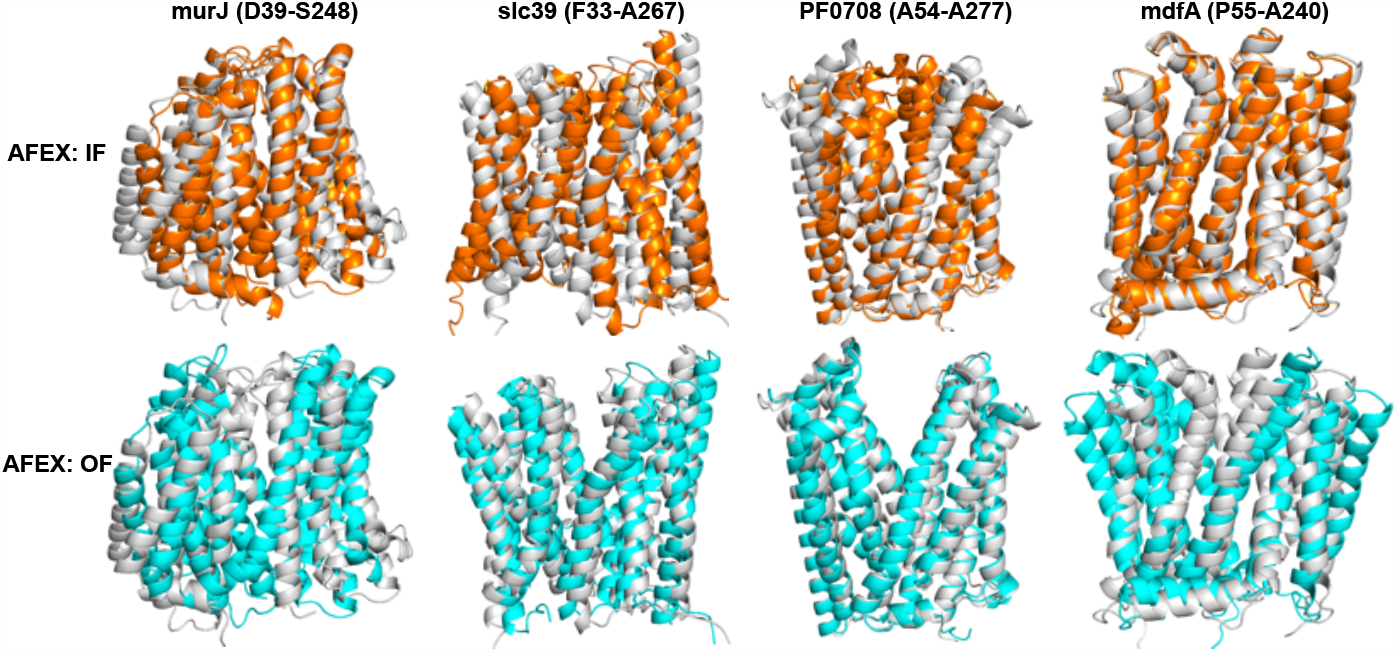
IF (orange) and OF (cyan) states generated by AFEX optimization, each aligned with the initial structure predicted by AF (grey). Protein names and selected gating residues are labeled.

### 4.3 AFEX generates inward-facing (IF) and outward-facing (OF) states of transporters

Obtaining alternative conformations for membrane transporters is notiously difficult for both experimental and computational techniques. This is easily achievable with AFEX, as demonstrated using four transporters from the IOMemP dataset [21]. Similarly, we define the CV loss as 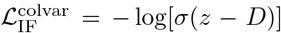 and 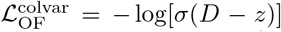, in which *D* is the C*α* distance between a pair of extracellular gating residues and *z* = 10 Å. With 500-step AFEX optimizations, conformational changes were induced towards either the IF or the OF states for all four proteins (Fig. 3). Take mdfA as an example: its initial structure with AF (grey) is in the IF state, and thus shows minimal change under AFEX conditioned on IF. However, when conditioned on OF, AFEX guides the protein into a OF conformational state. For PF0807, the initial AF structure corresponds to the OF state, and its IF state was successfully deduced by AFEX. Such a straightforward way to generate alternative states for membrane proteins will facilitate mechanistic studies of their functions.

## 5 Conclusion

In this study, we introduce AFEX as a versatile method that optimizes MSA embedding features under predefined conditions, guiding AF to explore conformational spaces for structures that meet these conditions. Using kinases and transporters as examples, we demonstrated that AFEX provides a controllable way for both the exploration and exploitation of conformational diversity. The condition, defined using CV loss, incorporates domain-specific insights. While simple, nearly binary variables were employed in this study, AFEX can condition on any local or global structure features that can be described by atomic coordinates. Adding scoring functions to the CV loss could further extend AFEX into tools for fully flexible protein-protein, peptide-protein, or ligand-protein docking.

